# Kinetics of winter deacclimation in response to temperature determines dormancy status and explains budbreak in different *Vitis* species

**DOI:** 10.1101/256362

**Authors:** Alisson P. Kovaleski, Bruce I. Reisch, Jason P. Londo

## Abstract

Bud dormancy and cold hardiness are critical adaptations for surviving winter cold stress for temperate perennial plant species, with shifting temperature-based responses during the winter. The objective of this study was to uncover the relationship between dormancy transition (chilling requirement) and temperature on the loss of cold hardiness and budbreak. Dormant cuttings of *Vitis vinifera*, *V. aestivalis*, *V. amurensis*, and *V. riparia* were examined to determine the relationship between chilling requirement and temperature on rate of deacclimation (*k*_deacc_). Differential thermal analysis was used to determine *k*_deacc_ using mean low temperature exotherms. Effect of chill was evaluated as the deacclimation potential (ψ_deacc_), which was the change in *k*_deacc_ due to chill accumulation. Budbreak was also evaluated in fully chilled buds at different temperatures. Results indicate that ψ_deacc_ varies dependent on dormancy state, following a logarithmic response to chill accumulation. The effect of temperature on *k*_deacc_ was exponential at low and logarithmic at high temperatures. The combination of ψ_deacc_ and *k*_deacc_ resulted in good prediction of deacclimation. Budbreak phenology was also explained by differences in *k*_deacc_. Deacclimation rates can be used as a quantitative determinant of dormancy transition and budbreak, and to refine models predicting effects of climate change.

## 4. Introduction

Due to their general stationary habit, plants have evolved many different coping mechanisms to survive stressful environments such as drought and temperature. With regard to winter in temperate climates, annual plants typically survive as dehydrated seeds while perennial plants develop dormancy and cold hardiness. Dormancy is the temporary cessation of visible growth in meristem containing structures, such as buds, and is divided in three types: para-, endo-, and ecodormancy (Lang, Early, Martin, & Darnell, 1987). Paradormancy is the suspension due to physiological factors within the plant but outside of the dormant structure, such as the suppression of lateral growth by the apical meristem (i.e. apical dominance). Paradormant buds transition into endodormancy as daylength and temperatures decrease, a state suspension of growth due to unknown endogenous factors within the dormant structure, preventing growth during times when environmental conditions fluctuate between conducive and inhibitory (Horvath, Anderson, Chao, & Foley, 2003). During endodormancy, it is generally understood that exposure to low, non-freezing temperatures is necessary (chilling requirement) to transition to the third dormancy state, ecodormancy. Tissues maintain a dormant state during ecodormancy due to unsuitable environmental conditions.

Concomitantly with the onset of dormancy, plants from areas where temperatures drop below freezing develop cold hardiness through different mechanisms. Some plants, or plant tissues, tolerate intracellular ice formation, in which cells are extremely dehydrated by the formation of extracellular ice, and the remaining water is bound to proteins (Burke, Gusta, Quamme, Weiser, & Li, 1976) likely solidifying in a glass-like state. Other plants present a freezing resistance mechanism, where water supercools in the intracellular spaces (Burke et al., 1976). Supercooling is a process through which water can remain in liquid state below its equilibrium freezing point, down to a maximum of ~ -40 °C (Bigg, 1953). These plants are typical of mid-latitude temperate climates, where temperatures do not drop below -40 °C during the winter. In buds that present supercooling ability, such as those of grapevines (*Vitis* spp.), cold hardiness may be measured through differential thermal analysis (DTA). This method uses thermoelectric modules that record voltage changes associated with the release of heat due to phase change of water into ice (Mills, Ferguson, & Keller, 2006). In grapevine buds, two peaks can be identified using DTA: a high temperature exotherm (HTE), typically around – 5 °C; and a low temperature exotherm (LTE), which varies depending on the conditions experienced by the bud. The HTE represents the freezing of extracellular water (Andrews, Sandidge, & Toyama, 1984), which is normal to the process of resisting cold. The LTE, however, is the freezing of intracellular water, and it happens when the supercooling point has been exceeded (Mills et al., 2006). The LTE represents the lethal temperature for a given bud, is correlated with manual assessments of bud death (Wolf & Cook, 1994), and mean LTE has been adopted as a consistent measurement of bud cold hardiness.

A general U-shaped pattern of changing cold hardiness is followed during the winter through three different stages: acclimation, maintenance, and deacclimation. This pattern is thought to be primarily driven by air temperature (Ferguson, Tarara, Mills, Grove & Keller, 2011; Ferguson, Moyer, Mills, Hoogenboom, & Keller, 2014; Londo & Kovaleski, 2017), although other climatic aspects may also be important (Antivilo et al., 2017). Acclimation appears to occur primarily during endodormancy, while deacclimation is enhanced during ecodormancy (Ferguson et al., 2011, 2014; Londo & Kovaleski, 2017). Depending on the climate and genotype, the transition between endo- and ecodormancy may occur toward the end of the acclimation stage, or during the maintenance stage of cold hardiness. This transition is typically evaluated by collecting dormant buds from the field and placing them in growth permissive conditions, followed by monitoring the time needed to reach budbreak (Weinbaum, Polito, & Muraoka, 1989; Lloyd & Firth, 1990; Cook & Jacobs, 2000; Fan et al., 2010; Zhang & Taylor, 2011). The chilling requirement has been met when 50% budbreak occurs within a time threshold {e.g. 3 weeks for peaches [*Prunus persica* (L.) Batsch; Lloyd & Firth, 1990]; 4 weeks for grapevines (Londo & Johnson, 2014)}. This method represents a subjective threshold for the change in endo-to ecodormancy transition. As a result, determining the molecular and metabolic cues and the consequences of this transition, such as budbreak, are very difficult to determine and poorly understood (Penfield, 2008).

As the dynamics of acclimation and deacclimation change with dormancy state (Ferguson et al., 2011, 2014), and dormancy state is determined by accumulation of chilling hours, a future climate may create issues for both cold hardiness and budbreak phenology. Average global temperatures are increasing (Walsh et al., 2014), although episodes of acute cold weather in the northern hemisphere are likely to increase as well through more frequent arctic oscillations (Kolstad, Breiteig, & Scaife, 2010). In addition to direct effects to cold hardiness, increasing temperatures will enhance chill accumulation in higher latitudes, while lower latitudes will experience a decrease (Luedeling, Girvetz, Semenov, & Brown, 2011). Changes in chill accumulation will modify the dynamics of dormancy transitions both in deciduous tree crops and other woody perennials, causing excessive responsiveness to warm spells or erratic budbreak and growth. This will ultimately require a shift towards higher latitudes and elevations of tree crops and assisted migration of forest species as they lag behind their optimal climate niche (Gray & Hamann, 2013). Therefore, it is important to understand how both temperature and dormancy status affect the loss of cold hardiness. The objective of this study was to better predict and understand the relationship between chill accumulation, the dormancy transition, and the loss of cold hardiness in wild and cultivated grapevines.

## 5. Materials and methods

Buds of four different species were collected multiple times during the dormant seasons of 2014/15, 2015/16, and 2016/17. *V. vinifera* L. ‘Cabernet Franc’, ‘Cabernet Sauvignon’, ‘Riesling’, and ‘Sauvignon blanc’ were collected from local vineyards (42.705N, 76.973W; and 42.845N, 77.004W), while *V. aestivalis* Michx. (two clones: PI483138, PI483143), *V. amurensis* Rupr. (three clones: PI588632, PI588635, PI588641), and *V. riparia* Michx. (four clones: PI588275, PI588562, PI588653, PI588711) were collected from the USDA Plant Genetic Resources Unit in Geneva, New York. For *V. vinifera*, buds from nodes 3 to 20 from the base of a cane were used, while for the other species, due to constraints regarding the number of clones, buds beyond position 20 were used as in Londo & Kovaleski (2017).

Hourly weather data from the closest Network for Environment and Weather Applications station (NEWA; http://www.newa.cornell.edu/) was used to compute chill accumulation using the “North Carolina” model (Shaltout & Unrath, 1983). The start date for each season was chosen based on when chill started to consistently accumulate instead of being negated, as determined by the NC model. These dates were 11 Sept 2014, 19 Sept 2015, and 24 Sept 2016.

Upon collection, canes were prepared into single-or two-node cuttings and placed submerging the basipetal cut surface in cups of water. The cups were placed into growth chambers at 2, 4, 7, 8, 10, 11, 22, or 30 °C. Not every genotype or temperature was used at all collection points, but this information is available in Supplementary Table S1. Differential thermal analysis (DTA) was performed to determine the cold hardiness of individual buds as their low temperature exotherm [LTE; see Mills et al. (2006) for details]. Briefly, buds are excised from the cane and placed on a thermoelectric module in a plate. These plates are placed in a programmable freezer with a cooling rate of -4 °C hour^-1^. Changes in voltage due to heat release in the freezing of water are measured by the thermoelectric module and recorded using a Keithley data logger (Tektronix, Beaverton, OR) attached to a computer. Between 4 and 8 buds were used at any time point to determine mean LTE, and the intervals between measurement days varied between chill accumulation level and temperature treatment, with details provided in Supplemental Table 1. Because lower rates of deacclimation were expected in lower temperatures and lower chill accumulations, lower temperature treatments were typically surveyed with wider separation between time points compared with high temperature treatments (Table S1).

In addition, an experiment was designed to compare temperature effects on fully chilled grapevines (1440 chill accumulation). Buds held at constant temperatures (2, 4, 7, 11, 22 °C) had their budbreak recorded following the modified E-L scale (Dry & Coombe, 2004) for all *V. vinifera* and *V. riparia*. For this, 5 buds were randomly selected and E-L number recorded in the same sampling interval as for LTE (Table S1) until buds were past stage 3 or bud material was exhausted.

### Statistics

Based on sample size across temperatures and across sampled years, data sets for *V. vinifera* were analyzed separately for each cultivar, while data for accessions of *V. aestivalis*, *V. amurensis* and *V. riparia* were combined at the species level. Analyses were separated into the effect of chill accumulation and the effect of temperature on rate of deacclimation. The datasets used for each analysis are specified in Supplementary Table S1.

#### Effect of Chill Accumulation

In order to assess the effect of chill accumulation in the rate of deacclimation (*k*_deacc_), or the deacclimation potential (ψ_deacc_), individual rates were calculated using linear regression in R (ver. 3.3.0, R Foundation for Statistical Computing) for each temperature and chill accumulation as factors. Although every temperature within a chill accumulation had the same data for day 0 (field collection), temperatures were still allowed to have different intercepts in order to reduce the effect of day 0 on *k*_deacc_. From this regression model, data points that had a studentized residual ≥2.5 were considered outliers and removed from the dataset, and the model was re-fit. The *k*_deacc_ at each chill accumulation were then transformed to percentage for 4, 7, 10, 11, and 22 °C, standardizing to their *k*_deacc_ at highest chill accumulation (either 1440 – 7 and 11 °C – or 1580 chill units – 4, 10, and 22 °C). The ψ_deacc_ was estimated as a logistic regression. Initial estimation used the drc library, but final estimation used the nls() function with the port algorithm, following the equation:

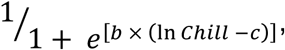 where *b* and *c* are the estimated parameters, and *Chill* is the chill accumulation at the time of sample collection. The parameter *c* is the inflection point of the logistic and *b* is the slope associated with the logistic regression.

#### Effect of Temperature

Initially, to study the effect of temperature, *k*_deacc_ obtained from multiple linear regression (temperature and chill accumulation as factors; Supplementary Table S1) in lower chill accumulations were re-scaled by dividing by the respective ψ_deacc_ (e.g., because the ψ_deacc_ was estimated to be at ~30% at 860 chill units for ‘Riesling’, the rates obtained from that data set were divided by 0.30). Since the effect of chill accumulation was estimated to be ~ 0 % for 360 chill units, these were excluded from the dataset for estimation of temperature effect. Arrhenius plots were created to evaluate temperature responses and data were then divided into two data sets, low (2 – 11 °C) and high temperatures (10 – 30 °C), and temperature effects on deacclimation rates were evaluated separately. The rationale for doing this is discussed below.

Instead of using the initially calculated rates derived from the above linear models, the complete data sets were used with the effect of temperature as a continuous variable. The values for the different chill accumulation points were normalized to that of full chill by multiplying the ψ_deacc_ at any given chill accumulation, effectively removing the effect of chill by mathematically reducing the time for deacclimation. Lower chill accumulations were thus included in the calculations instead of using only those where chill requirements were initially assumed to be fulfilled (1440 and 1580 chill accumulations).

For low temperatures, the effect was assumed to be an exponential, and therefore the rates were calculated as *k*_deacc_ = *m* × *e*^(*n* × *T*)^, where *m* and *n* are the parameters estimated, and *T* is the temperature in °C. For high temperatures, a logarithmic curve was used: *k*_deacc_ = *q* × ln(*T* – *r*), where *q* and *r* are the parameters estimated, and *T* is the temperature in °C. For both low and high temperatures, parameters were estimated using nls(), with the port algorithm.

#### Fitness of Effects of Chill and Temperature

The fits of all non-linear curves (effect of chill accumulation, low and high temperatures) were tested using Effron’s pseudo-R^2^ with the Rsq function in the soilphysics library, which is calculated using the sum of squares of ordinary residuals from the models. The final model for loss of cold hardiness from an initial point is then *Δ*_*LTE*_(*T, Chill*) = *k*_deacc_ × ψ_deacc_ × *t*, where the change in LTE (*Δ_LTE_*) is a function of the temperature (*T*) it was exposed to, how much chill accumulation (*Chill*) had passed before exposed to deacclimation temperatures, and the time (*t*) of exposure. A linear model was used to evaluate the accuracy of the predictions using the complete data set. For this, instead of temperatures, *k*_deacc_ = 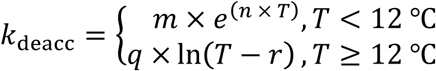 was used, and instead of accumulated chill, the ψ_deacc_ was used. From this model (ΔLTE = β × *k*_deacc_ × ψ_deacc_ × time), considering estimations were appropriate, the β associated with the estimated *k*_deacc_ and ψ_deacc_ was expected to be 1.

#### Budbreak

In order to evaluate whether the same temperature effects governing deacclimation and loss of cold hardiness (measured as LTE) also controlled differences in the temporal rate of budbreak, a linear model was used. For this, forward selection was conducted with a reduced model containing an intercept only, and a full model containing all possible interactions of growing degree-days [GDD = *k*_deacc_ × *t* (day)], temperature, species, and genotype as explanatory variables for bud growth stage. The forward selection was corrected based on the Bayesian information criterion (BIC).

## 6. Results

After removing outliers using multiple linear regression, data sets kept ≥ 90% of the observations for all genotypes, with the exception of *V. amurensis*, which had 19.5% of observations removed (Table 1). Linear behavior appears to describe well the effects of temperature and chill accumulation in the deacclimation of grapevine buds (Figure 1). A logistic regression appeared to be a good fit for the ψ_deacc_ (Figure 2), with pseudo-R^2^ values greater than 0.87 for all genotypes except for ‘Cabernet Sauvignon’ and *V. amurensis* (0.65 and 0.79, respectively; Table 1). The parameter *e* is the natural logarithm of the inflection point, and there was little difference in this parameter for these genotypes (e.g., 6.77 to 6.88, for ‘Sauvignon blanc’ and ‘Riesling’, respectively, which is equivalent to 872 to 972 chill units).

**Table 1.**
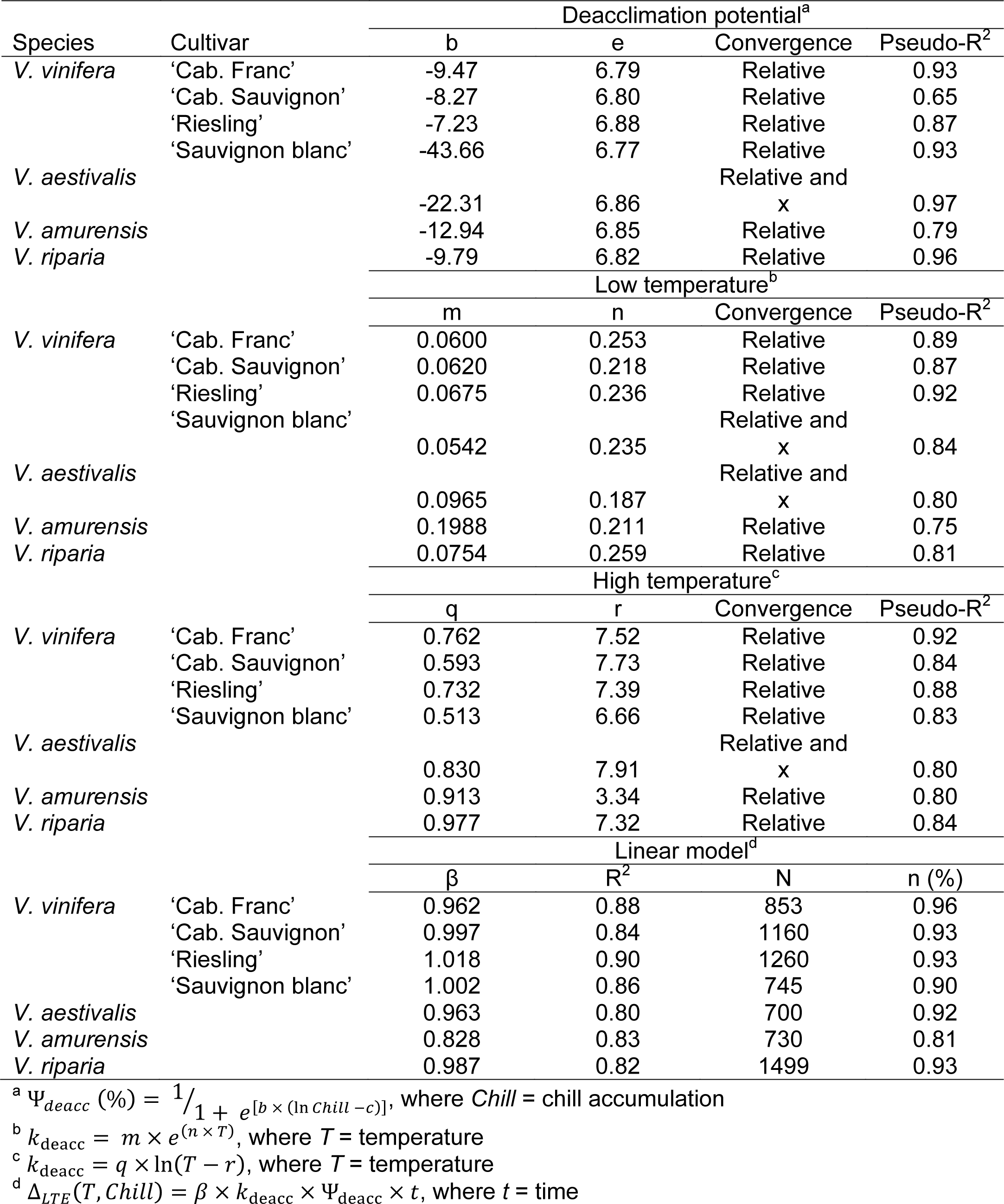
Parameter estimates and model fits for deacclimation potential, low and high temperature deacclimation rates, and linear relationship between LTE and the deacclimation rate × deacclimation potential for 7 *Vitis* genotypes.

**Figure 1.**
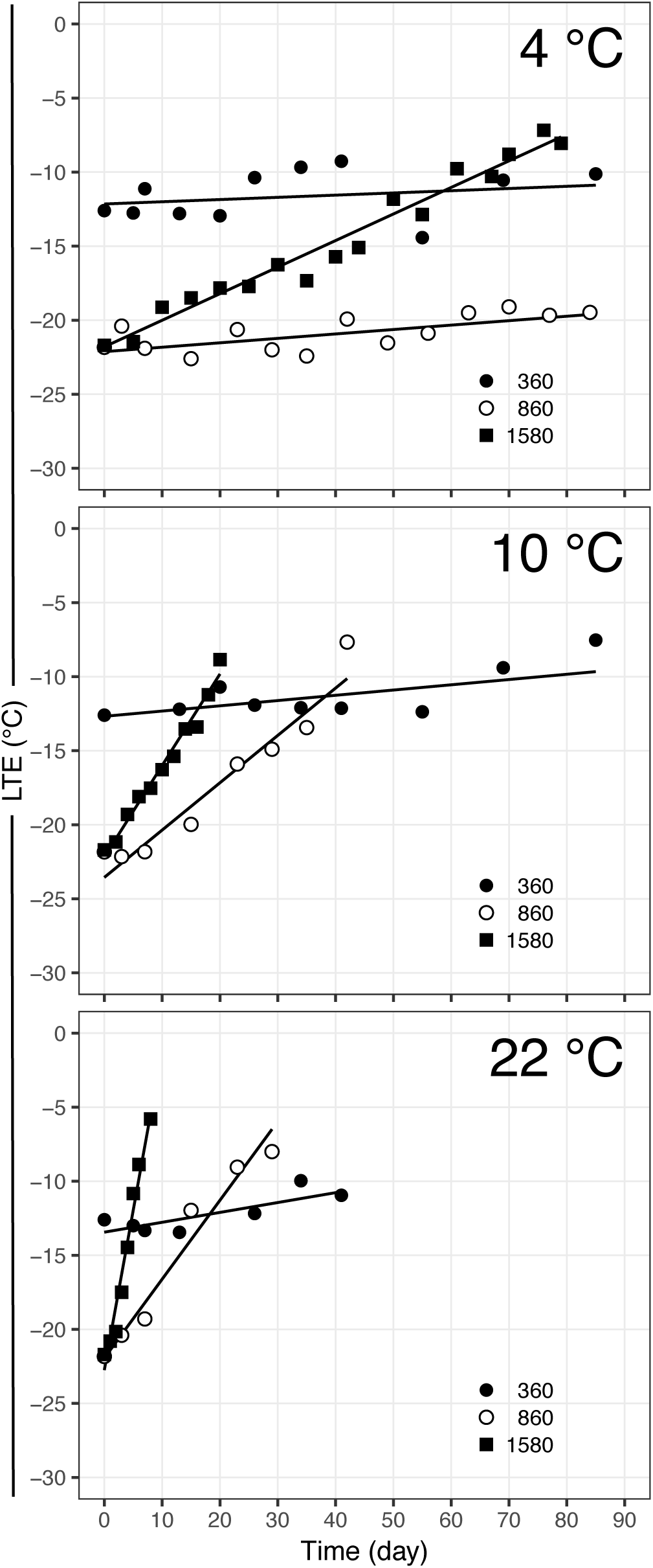
Deacclimation of *V. vinifera* ‘Riesling’ buds at three temperatures (4, 10, and 22 °C) collected from the field at three different chill accumulations (360, 860, 1580). *Buds at 360 chill accumulation were deacclimated at 11 °C.

**Figure 2.**
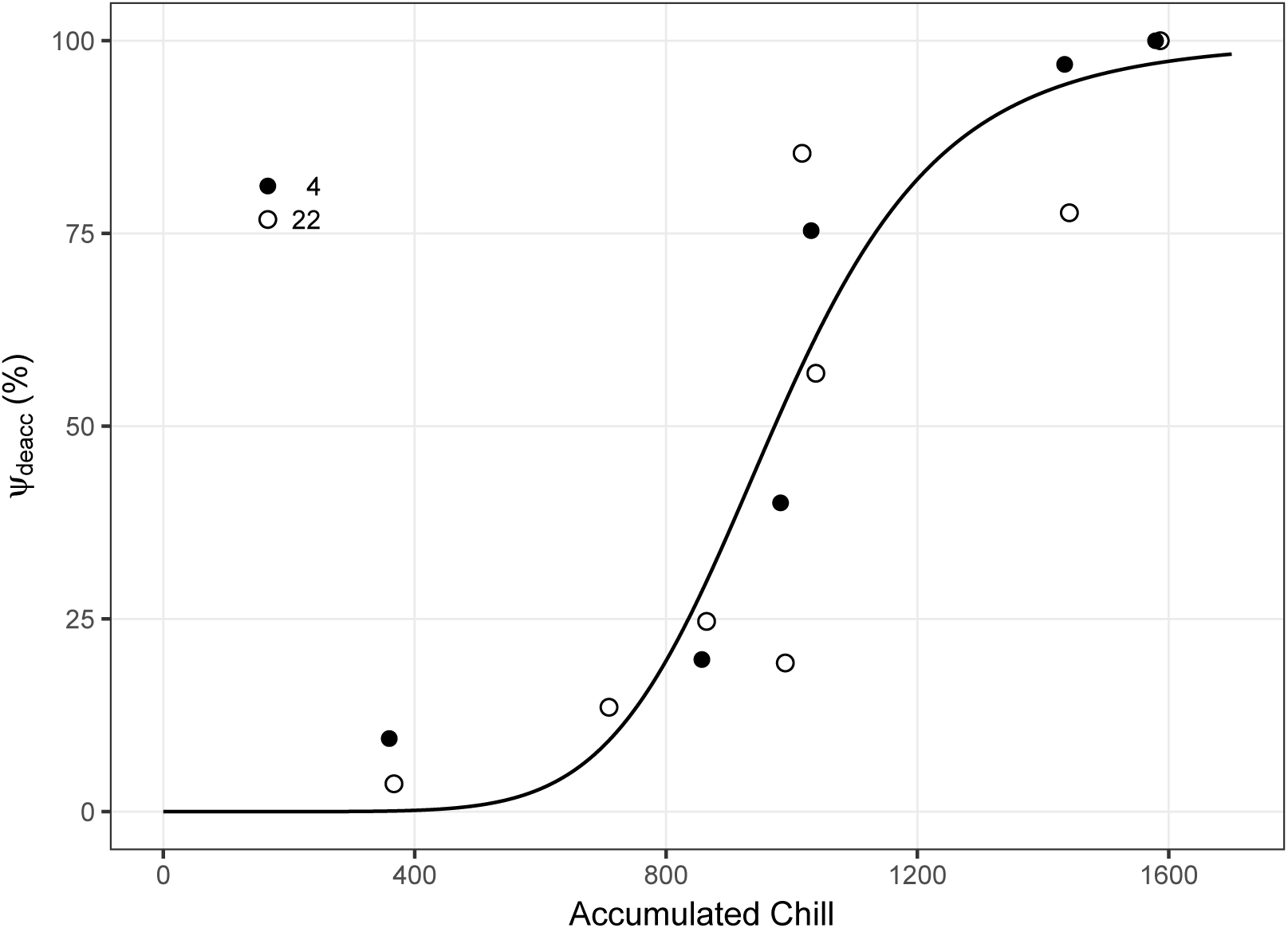
Deacclimation potential of *V. vinifera* ‘Riesling’ buds collected at different chill accumulations in two temperatures. Deacclimation rate at 1580 chill units was used as reference (100 %). The relationship between deacclimation potential and accumulated chill was described by the equation: Ψ_*deacc*_ (%) = 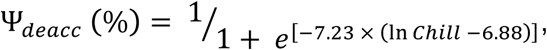 with a pseudo-R^2^ = 0.87.

Scaled deacclimation rates based on ψ_deacc_ and corresponding Arrhenius plot (Figure 3) demonstrated a clear discontinuity between low and high temperatures. A deceleration of the increase of *k*_deacc_ as temperature increased was observed, which justifies the different behaviors used (exponential and logarithmic for low and high temperatures, respectively). The estimation of temperature effects for low and high temperature intersect between 10 and 11 °C because of the uncertainty in the exact temperature where there is a change in behaviors. This is partially due to the lack of data points between 11 and 22 °C, although at 10 and 11 °C the calculated rates through either model would be very similar. All the *V. vinifera* cultivars had similar deacclimation rates at low temperatures, but ‘Riesling’ and ‘Cabernet Franc’ had higher deacclimation rates at high temperatures than ‘Cabernet Sauvignon’ and ‘Sauvignon blanc’ (Table 1, Figure 4). *V. riparia* and *V. aestivalis* had similar rates to those of *V. vinifera* at low temperatures, but *V. riparia* had estimated rates similar to *V. amurensis* in higher temperatures. *V. aestivalis* was similar to the faster *V. vinifera* (’Cabernet Franc’ and ‘Riesling’). *V. amurensis* showed the highest deacclimation rates at both low and high temperatures.

**Figure 3.**
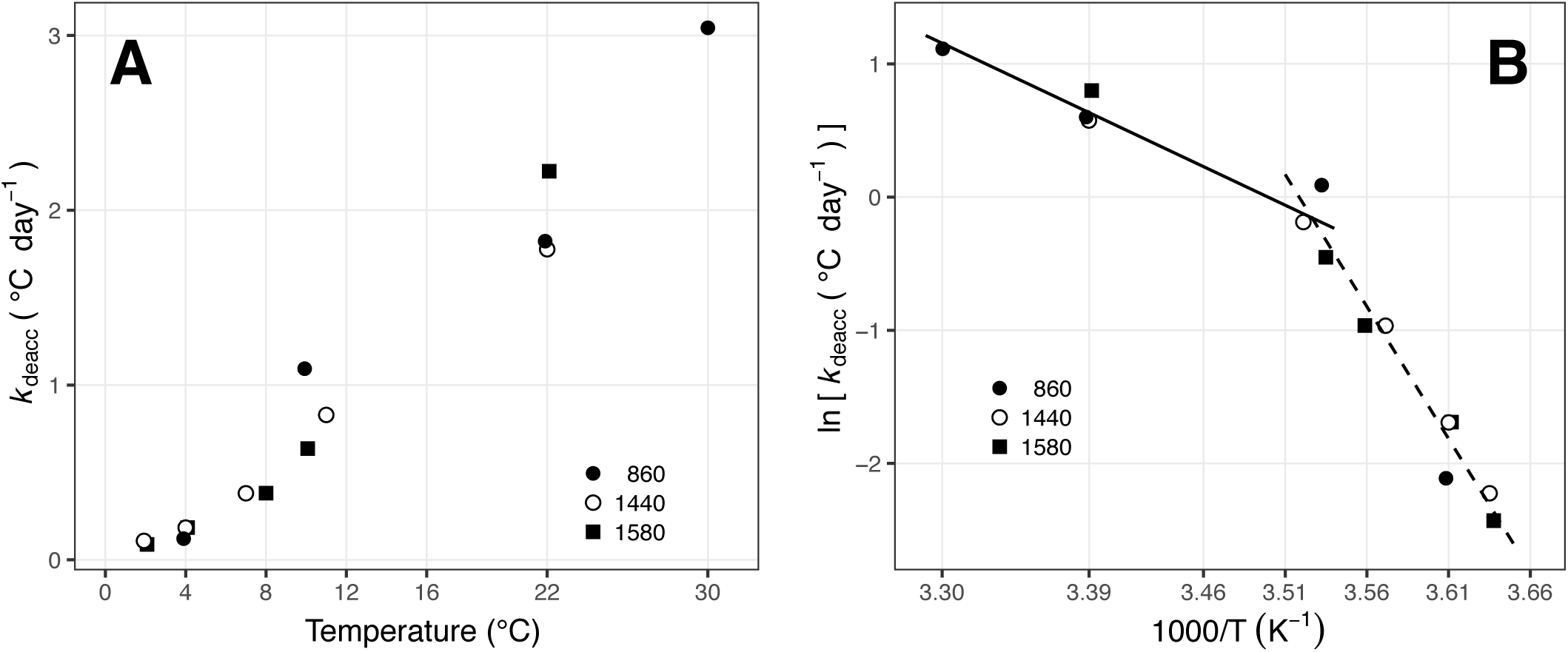
Effect of temperature on deacclimation rates of *V. vinifera* ‘Riesling’. A) Scaled deacclimation rates (based on deacclimation potential) at three different chill accumulations in response to temperature; B) Arrhenius plot of scaled deacclimation rates. Tick marks in both panels are equivalent and mirrored (e.g. 0 °C = 3.66 × 1000 K^-1^).

**Figure 4.**
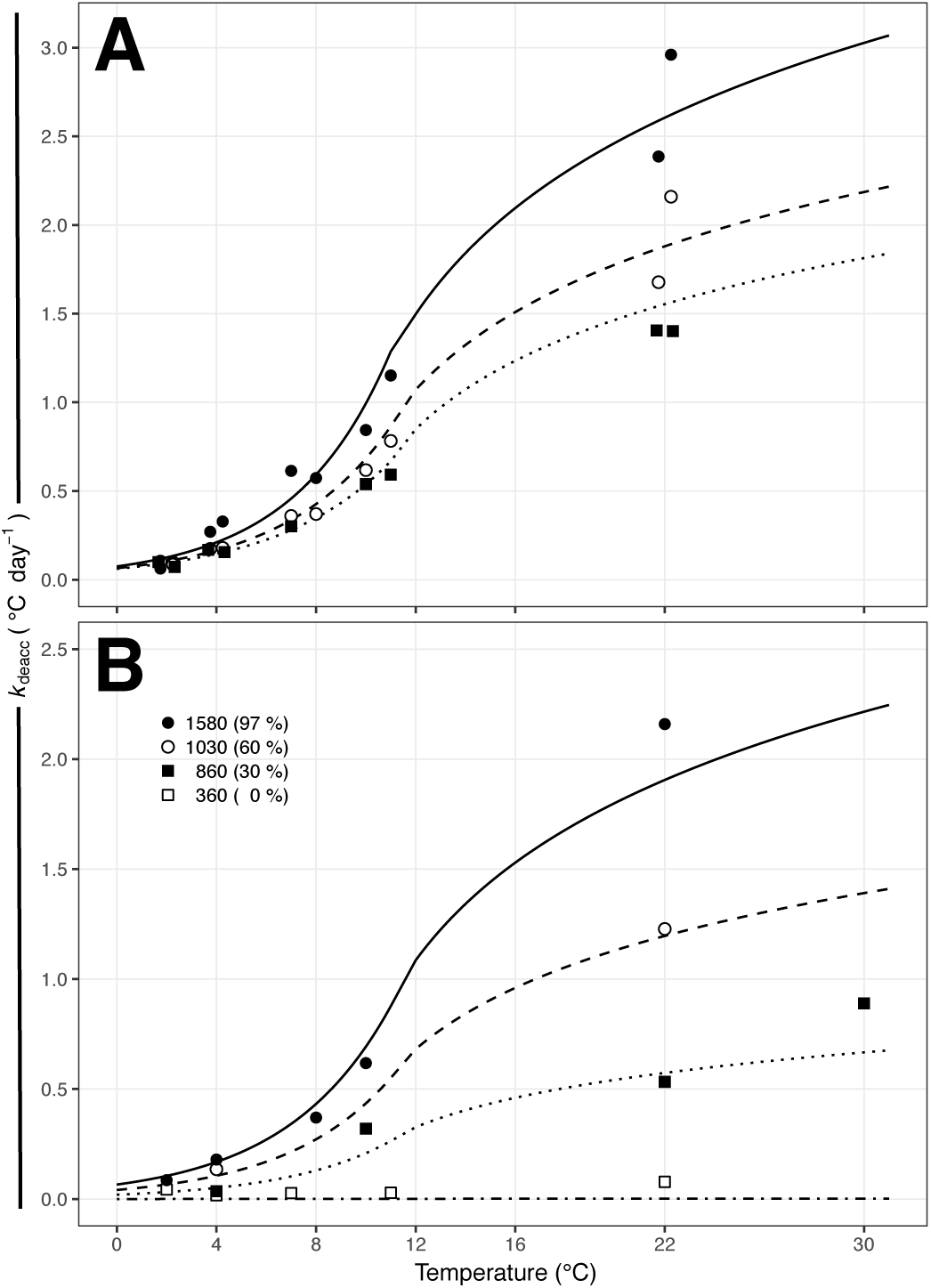
Deacclimation rates of *Vitis* buds as a function of temperature. A) Deacclimation rates at >90 % deacclimation potential for three different *Vitis* genotypes: *V. riparia* (closed circle - full line), *V. vinifera* ‘Riesling’ (open circles - dashed line), and *V. vinifera* ‘Cabernet Sauvignon’ (closed squares - dotted line); B) Deacclimation rates of *V. vinifera* ‘Riesling’ at different chill accumulations as a function of temperature: 1580 (closed circle – full line), 1030 (open circle – dashed line), 860 (closed square – dotted line), 360 (open square – dashdotted line).

Linear models showed a good fit for all genotypes (R^2^ ≥ 0.80). *V. vinifera* cultivars, which were analyzed separately had higher R^2^ values. The better fit for V. *vinifera* cultivars is also seen in their β coefficients, which are very close to 1, although *V. aestivalis* and *V. riparia* had βs of 0.99 and 0.96 respectively. *V. amurensis* had the lowest β, at 0.83.

The final model in the multiple linear regression for budbreak stage had a P<0.001 and adjusted-R^2^ = 0.66 using an intercept and GDD × species interaction *(Budbreak stage (species)* = *a + b* × *GDD*). The effect of *k*_deacc_, i.e. the effect of temperature, on budbreak can be seen in figure 5. When LTE and budbreak are regarded in respect to time, a clear separation was observed between temperatures (Figure 5A, C, E, G). When GDDs are used, accounting for differences in *k*_deacc_, LTE and budbreak stages for all temperatures are “stacked” (Figure 5B, D, F, H). Within the same temperature, budbreak occurs at different times for different genotypes (Figure 6A), but is grouped within species when using GDDs (Figure 6B).

**Figure 5.**
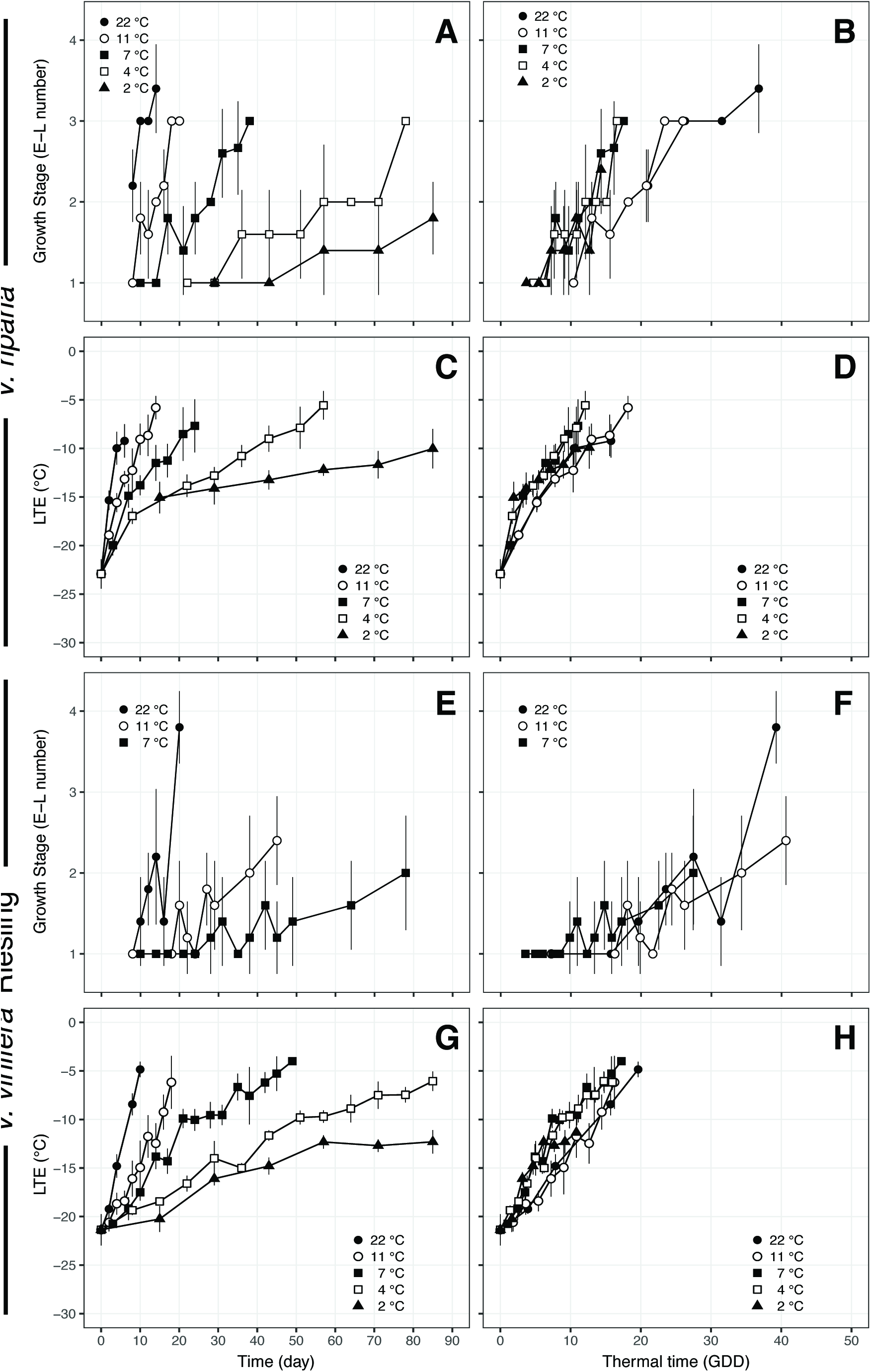
Deacclimation and bud phenology of *V. riparia* and *V. vinifera* ‘Riesling’ in relation to time (A, C, E, G) and thermal time in GDD (B, D, F, H). Bud phenology as the “E-L number” (Dry and Coombe, 2004) is shown for 5 temperatures in *V. riparia* and 3 temperatures in ‘Riesling’, as no bud development was recorded within 90 days at 2 and 4 °C for ‘Riesling’. GDDs were calculated based on *k*_deacc_ for each genotype (see Table 1) at any given temperature multiplied by time in days.

**Figure 6.**
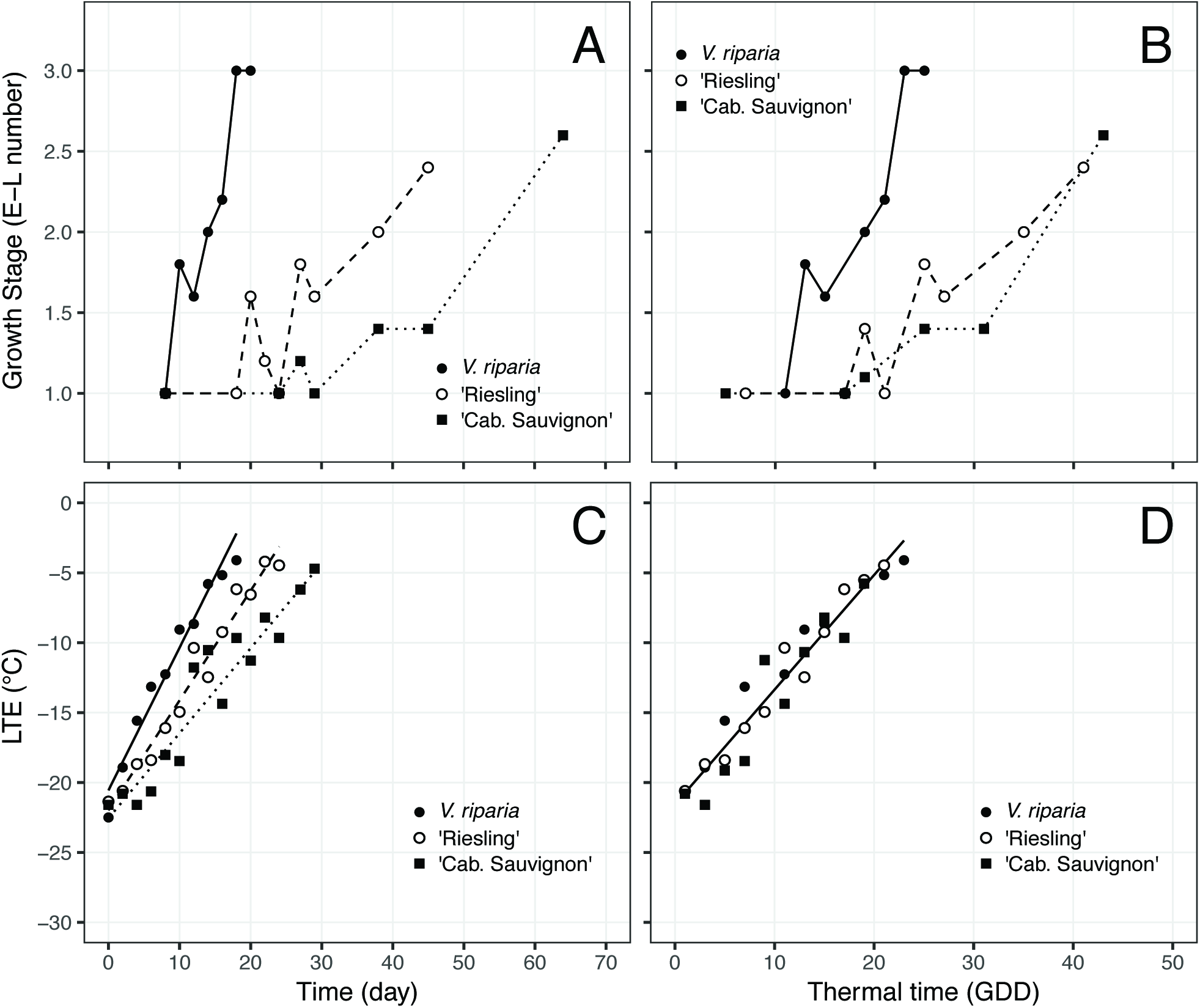
Deacclimation and bud phenology of *V. riparia* (closed circle – full line), *V. vinifera* ‘Riesling’ (open circle – dashed line), and *V. vinifera* ‘Cabernet Sauvignon’ (closed square – dotted line) at 11 °C in relation to time (A, C) and thermal time in GDD (B, D).

## 7. Discussion

Chilling accumulation, dormancy state, cold hardiness and budbreak are complex traits driven by the interaction of physiological and climactic attributes. The consistency of winter temperatures is expected to be a major aspect impacting the sustainability and survival of perennial species in a future climate. This study was conducted to gain a better method for predicting dormancy transition and chilling requirement in grapevine, as well as to understand how deacclimation processes (loss of cold hardiness) and resulting budbreak are impacted by dormancy state.

This paper provides a quantitative assessment of the transition from endo-to ecodormancy, as evaluated by the ψ_deacc_. Unlike the typical dormancy assessment that uses time to budbreak as a measurement of the transition (Londo & Johnson, 2014), using the deacclimation rates does not require subjective assumptions currently used to determine chilling fulfillment; such as having 50% budbreak occur within 28 days. Budbreak occurs only after the freeze mechanisms associated with cold hardiness are lost (Figure 5, 6; Ferguson et al., 2014; Salazar-Gutiérrez, Chaves, Anothai, Whiting, & Hoogenboom, 2014). Thus, the amount of time needed to reach 50% budbreak in forcing assays is a combination of the time needed to fully deacclimate, and time required for deacclimated buds to reach a visible change in phenological stage. This study leverages the ability to assess loss of hardiness through decreasing supercooling ability as a way to quantitatively assess rates of deacclimation and its relation to changes in chilling exposure.

Mid-winter warming events are of major concern for temperate crops, yet natural areas are also affected. In native trees within natural ranges, little to no damage occurs during the autumn because the freeze resistance readily withstands the minimum temperatures they usually encounter (Vitasse, Lenz, & Körner, 2014). However, damage can be substantial in late freeze events during the spring or unusual warm events in early spring as the potential for deacclimation increases exponentially (Vitasse et al., 2014). For example, a late spring freeze in 2007 in North America caused damage in native forests, delaying and reducing canopy establishment by 16-34 days (Gu et al., 2008). Bokhorst, Bjerke, Tømmervik, Callaghan, & Phoenix (2009) reported that dwarf shrub vegetation in a sub-Artic region where midwinter temperatures rose to ~7 °C for 2 weeks followed by further cold temperatures resulted in an 87% reduction in growth during the summer compared to neighboring undamaged areas. Cannell & Smith (1983) show results from different authors for forest species in which the time to budbreak is reduced in an exponential manner. This is similar to our assessment of ψ_deacc_, where the upper portion of the logistic is like the exponential decrease in time to budbreak. The logistic behavior of ψ_deacc_ in relation to chill accumulation explains the continuous increase risk of damage of mid-winter warm spells the later they occur until 100% ψ_deacc_ is achieved. This is in contrast with the model developed by Ferguson et al. (2011; 2014), where a hard limit for the dormancy boundary is used to model grape cold hardiness. It is likely that, similar to the behavior described in our study, forest species also have increased deacclimation rates with higher chill accumulation, seen as a decrease in the time to budbreak. Additionally, it is important to note that while canopies may be able to recover from midwinter bud kill events, damage to the reproductive buds in the case of fruit and nut crops will reduce yields in the season following damage.

Our assessment of ψ_deacc_ leading to increases in *k*_deacc_ agrees with the general observation that warm temperatures in the early winter are negligible for bud growth (Cannell & Willett, 1975) and that increasing chill accumulation reduces the heat requirements for budbreak in fruit species (Couvillon & Erez, 1985; Citadin, Raseira, Herter, & Silva, 2001), and forest species (Hunter & Lechowicz, 1992; Dantec et al., 2014), considering that loss of hardiness is required for budbreak to occur (Figure 5, 6; Ferguson et al., 2014). Analyzing bud phenology at approximately full chilling (1440 Chill accumulation ≥ 95% ψ_deacc_ for all genotypes), we have seen that *k*_deacc_ is responsible for differences in early phenological development in the spring between genotypes within *V. vinifera*, but species still had different phenology with regard to GDD. Planted in the same environment, Taulavuori et al. (2004) observed that deacclimation occurs faster in mountain birch [*Betula pubescens* ssp. *czerepanovii* (Orl.) Hämet-Ahti] ecotypes with origin in areas of low chill accumulation. If the mechanisms are similar to those described here, what may have been observed in their study was not different rates of deacclimation, but plants that were at different stages in their deacclimation potential, or a combination of both. Climate change appears to speed the loss of cold hardiness and promote earlier leaf-out, although the response is not linear due to interactions with chilling and photoperiod (Vitasse et al., 2014). In our case, the predicted increase in chill accumulation (Luedeling et al., 2011) will result in higher ψ_deacc_ and higher risk of damage should midwinter warm spells increase in a future climate (Vitasse et al., 2014; Walsh et al., 2014). However, earlier budbreak may not lead to higher frequency of damage due to frost events if consistently higher spring temperatures (Chmielewski, Götz, Weber, & Moryson, 2018) accompany future climate changes.

Pagter & Arora (2013) suggest that deacclimation rates are not linear, and that a lag phase may occur during initial exposure to warm temperatures. The initial lag phase might explain the differences in deacclimation rates measured in Cragin, Serpe, Keller, & Shellie (2017) compared to those reported in this study, as they only evaluated deacclimation for 4 days. This may also suggest why they had differences in rates across the years, although this may also be an effect of different chill accumulation between the two years in their study. Ferguson et al. (2011; 2014) used asymptotic bounds added through a logistic component, deviating from a strictly linear rate. In our data, although in some cases an initial curvature of the data points can be observed (e.g., Figure 1, 10 °C – 860 chill units), linear rates were strong descriptors of the deacclimation rates as observed in the fits of our calculated rates of deacclimation and in the early multiple linear regression models with temperature and chill accumulation as factors (R^2^ > 0.83, p < 0.001 for all cultivars; Table S1), similar to Wolf & Cook (1992), also looking at deacclimation rates for grapevines – although they only used 1 temperature. We also stopped measuring LTEs when those were close to -5 °C, where we had difficulties reading LTE peaks due to the merge with the much larger high temperature exotherms, and therefore any curvature in the higher portions of the deacclimation curve would not be observed. Such curvature is observed in Arora et al. (2004) and Kalberer, Leyva-Estrada, Krebs, & Arora (2007), although hardiness loss stopped at a lower temperature, which was speculated to be due to lack of chill.

Ferguson et al. (2011; 2014) used a linear response of the rates of deacclimation in relation to temperature for 21 *V. vinifera* and two *V. labruscana* Bailey cultivars, with a different rate and base temperature for endo-and ecodormant buds. In our experiments, deacclimation occurred in temperatures below those established as the threshold for deacclimation by Ferguson et al. (2014). This suggests that although it may produce a good fit for field data, the current model may be over-simplifying the response to temperature from a physiological standpoint. Takeuchi & Kasuga (2017) have also seen deacclimation through electrolyte leakage of bark and xylem of Japanese white birch [*Betula platyphylla* Sukatchiev var. *japonica* (Miq.)] at low temperatures, including high below-freezing temperatures (-5 to 0°C). In our study the temperature dependency of *k*_deacc_ appears to follow simple enzymatic kinetics, affected by both the catalytic rate (Arrhenius’ law) in lower temperatures, and the denaturation of enzymes in higher temperatures. This behavior suggests that an active temperature sensing, as defined by Ruelland & Zachowski (2010), is not necessary for deacclimation. Although deacclimation is likely controlled by a complex metabolic chain or cycle, the rates of any given process as a whole are defined by the slowest enzyme – which may change depending on the temperature (Ruelland & Zachowski, 2010). Additionally, Farquhar & Richards (1984) proposed that whole plant processes should be analyzed as simple chemical processes in respect to temperature. Therefore, further comparisons are made in this discussion with activities of single enzymes and whole metabolic cycles without distinction.

It is notable in the deacclimation process measured in this study that rates are departing from the exponential increase due to temperature in low range (estimated as ≥ 12 °C) as seen in the Arrhenius plot (Fig. 3B). This would suggest that some limitation occurs at higher temperature to the process as a whole: either substrate limitations or enzyme denaturation (Truhlar & Kohen, 2001). It is important to note, however, that there is no drop in the *k*_deacc_ from 22 to 30 °C, but a reduction on the increase of *k*_deacc_ as compared to from 2 to 10 °C (Fig. 3). This differs from other processes in grapevines, such as root respiration, which has an exponential response to temperature up to 32 °C for short-term exposure (Huang, Lakso, & Eissenstat, 2005). This is likely an adaptation of the enzymes involved in the process of deacclimation, considering low temperatures in early spring when deacclimation occurs. Rigano et al. (2006) demonstrated similar kinetics when comparing the activity of methyl viologen-dependent nitrate reductase extracts from psychrophilic and mesophilic algae. Arrhenius plots of the psychrophilic alga had an inflection point at 15 °C, whereas the mesophilic alga had an inflection point at 25 °C. We made no attempt in our study to find different inflection points for each species, as we lacked data in the 12-21 °C range. However, future studies could be conducted to examine if deacclimation rate is adaptive in widely distributed *Vitis* species as well. The activity of some enzymes, such as catalases and peroxidases, appears to be correlated with release of dormancy (Farokhzad, Nobakht, Alahveran, Sarkhosh, & Mohseniazar, 2018) and subsequent budbreak (Pérez & Burgos, 2004). It may be interesting to measure the activity of these enzymes at different temperatures to evaluate whether they have a similar response to temperature as that evaluated in the deacclimation here. Other enzymes involved in the synthesis and degradation of plant hormones might also be involved, considering that growth inhibition by abscisic acid or promotion by gibberellins and cytokinins appear to interplay in the control of dormancy (Horvath et al., 2003).

In a review of photosynthesis responses to temperature, Berry & Björkman (1980) showed that the phase of exponential increase in photosynthesis rate can end at temperatures as low as 10 °C and as high as 40 °C, and that both the rates and the optimum temperatures are also affected by the temperature regime in which plants are grown. For deacclimation rates in grapevine buds, Cragin et al. (2017) suggested an effect of the field LTE (or the intercept) on the deacclimation rate, which would indicate an effect of the environment. We did not attempt to use field LTE as a factor in our calculations; however, we expect that it would not greatly impact *k*_deacc_: both 860 and 1580 chill units had similar field LTEs, but very different *k*_deacc_ (Fig. 1). We believe that most of the environmental effect on the *k*_deacc_ is well described by the chill accumulation through the ψ_deacc_.

Cultivar or accession differences are seen within species in cold hardiness (Arora et al., 2004; Ferguson et al., 2011, 2014; Salazar-Gutiérrez et al., 2014; Salazar-Gutiérrez, Chaves, & Hoogenboom, 2016; Szalay, Molnár, & Kovács, 2017), although deacclimation rates may or may not differ (Arora et al., 2004; Ferguson et al., 2011, 2014; Szalay et al., 2017). The effect of genotypes within species in *k*_deacc_ is suggested by the better fit of our curves to *V. vinifera* cultivars as compared to the other species, where parameters were estimated by combining different accessions in the wild material. However, our dataset did not allow us to properly analyze each genotype of the wild accessions separately. It has been suggested that there is no correlation among species between the maximum cold hardiness or climate of origin and the rate of deacclimation (Arora et al., 2004; Kalberer, Wisniewski, & Arora, 2006). Vitasse et al. (2014), however, suggests that the rate of deacclimation is more related to temperature fluctuations in the area where species evolved, diminishing the potential for deacclimation in locations where frequent temperature fluctuations occur in the winter. In our case, *V. riparia* and *V. amurensis*, two species from colder climates as compared to *V. vinifera*, had higher *k*_deacc_ at all (*V. amurensis*) or at moderate to high temperatures (*V. riparia*) than *V. vinifera. V. aestivalis*, however, did not appear to differ from the *V. vinifera* cultivars, in agreement with the assessment by Kalberer et al. (2006). Kalberer et al. (2006) also suggested that field and controlled studies can result in different responses to temperature. In our study, however, we observed higher rates of deacclimation in *V. amurensis* and *V. riparia* compared to *V. aestivalis*, which is comparable and follows the same behavior as the “responsiveness” described by Londo & Kovaleski (2017) using field observations.

Cannell & Smith (1983) speculate that average daily temperature is a good descriptor for timing of budbreak, and that daily amplitude and photoperiod may be ignored. Similarly, Antivilo et al. (2017) saw no or little differences in deacclimation due to low and high temperature amplitudes, respectively. Although this was not tested, it appears that a linear approximation may be used for temperature effects on deacclimation, especially in small temperature amplitudes. Therefore, using daily average temperatures may be appropriate for field data. However, long-term averages could result in larger error in assessments, and physiological-based studies should use more precise descriptions of temperature effects such as those reported here.

It is possible that the accuracy of models for prediction of budbreak may be increased if they are attached to predictions of cold hardiness and deacclimation kinetics. Rubio, Dantas, Bressan-Smith, & Pérez (2016) reported both LTE and time to 50% budbreak for *V. vinifera* ‘Thompson Seedless’, which appear to be positively correlated in the acclimation period. Although the authors did not report chill accumulation, it is clear that cold hardiness increases (more negative LTEs) and days to budbreak decreases once daily mean temperatures below 15 °C start consistently occurring (May – Southern hemisphere). In their case the time to budbreak appears to reach the limit of endodormancy [50% budbreak within 28 days as per Londo & Johnson (2014)] in early June in Chile, but continue dropping until their last evaluation in August. This is in accordance with our assessment that as chill accumulates, there is an increase in ψ_deacc_, leading to potentially faster budbreak. Because budbreak occurs following loss of hardiness, and because of the relationship between ψ_deacc_ (through chill accumulation) and *k*_deacc_ leading to budbreak, we hypothesize that studies that identify earlier or faster budbreak due to the use of hydrogen cyanamide (Pérez & Burgos, 2004; Pérez & Lira, 2005) would likely find faster rates of deacclimation as well.

Bud phenology timing appears to follow the same temperature response as deacclimation within species. Therefore, it is likely that loss of cold hardiness may actually be early stages of growth within the bud, and that the differences in bud development in regard to GDDs (Figure 6B) between species are due to different bud morphology. However, there have been no studies comparing early morphological development in buds of different *Vitis* species. In *V. vinifera*, however, Andreini, Viti, & Scalabrelli (2009) showed a difference in thermal time requirements between different genotypes grouped in late, intermediate and early budbreak. They used the same temperature response for all cultivars, however. This is similar to the assessment of budbreak by Londo & Johson (2014), in which ‘Riesling’ and ‘Cab. Sauvignon’ had different times to budbreak within the same chill accumulation. In our case, it is clear that if the same effect of temperature was used between cultivars, budbreak would demand different thermal time accumulations for ‘Riesling’ (early budbreak) and ‘Cab. Sauvignon’ (late budbreak; Figure 6A). However, when we account for the difference in the efficiency of the use of temperature ( e.g., *k*_*deacc*_*Riesling*__ > *k*_*deacc*_*Cab.Sauvignon*__ at any given temperature), both *V. vinifera* genotypes have the same GDD requirements for budbreak (Figure 6B). Therefore, forcing assays using budbreak phenology imply less efficient deacclimators, such as ‘Cabernet Sauvignon’, require higher chill accumulation. Our results indicate that high-chill requirement phenotypes may actually be a combination of higher chill requirement and low temperature efficiency (low *k*_deacc_).

Specific studies linking environmental factors to understand plant phenology in order to evaluate impacts of climate change are needed (Cleland, Chuine, Menzel, Mooney, & Schwartz, 2007; Chmielewski et al., 2018). In this study, we demonstrate an objective method for determining the dormancy transition that could prove useful in other perennial species that utilize supercooling for cold hardiness [e.g. peach, cherry (*Prunus* spp. L.), azalea (*Rhododendron* spp. L.), larch (*Larix kaempferi* Sarg.)] in order to understand how they might respond to the future shifts in chill accumulation and increase in average temperature as a consequence of climate change. Although measurements of cold hardiness may be different for each species, it appears that this phenotype can aid in understanding dormancy transitions and spring phenology of plants, as we have demonstrated that early bud phenology and deacclimation to cold are linked and therefore results based on budbreak alone are confounded by dynamics of the loss of hardiness. It is clear that simple assessments of deacclimation rates simply using “endo-” and “ecodormant” material is not appropriate, as there is a continuum of response in deacclimation rates as chill is accumulated. Additionally, although we made no attempt to evaluate different chill accumulation models, it is possible that using deacclimation rates may be a more appropriate way of defining chilling temperatures, considering the more quantitative nature of this analysis.

## 8. Acknowledgements

The work reported here was partially supported by the National Institute of Food and Agriculture, U.S. Department of Agriculture, through the Northeast Sustainable Agriculture Research and Education program under subaward number GNE16-130; and by CAPES, Coordenação de Aperfeiçoamento de Pessoal de Nível Superior, Brazil. The authors would like to acknowledge and thank the following people for contributing to the field collection and processing of dormant bud tissues used in this study: Kathleen Deys, Bill Srmack, John Keeton, Bob Martens, and Matheus Baseggio. The authors also thank Dr. Tim Martinson, Cornell University, for use of freezer facilities, and Anthony Road Wine Co. and Ravines Wine Cellars for access to *V. vinifera* samples. The authors have no conflict of interest to declare.

## Supporting information

Table S1. Information associated with each deacclimation experiment for each genotype used. Linear models were fit per genotype using chill accumulation and temperature as factors.

## References

Andrews P.K., Sandidge III C.R. & Toyama T.K. (1984). Deep supercooling of dormant and deacclimating *Vitis* buds. American Journal of Enology and Viticulture 35, 175–177.

Andreini L., Viti R. & Scalabrelli G. (2009). Study on the morphological evolution of budbreak in Vitis vinifera L. Vitis 4, 153–158.

Antivilo F.G., Paz R.C., Keller M., Borgo R., Tognetti J. & Juñent F.R. (2017). Macro-and microclimate conditions may alter grapevine deacclimation: variation in thermal amplitude in two contrasting wine regions from North and South America. International Journal of Biometeorology 61, 2033–2045.

Arora R., Rowland L.J., Ogden E.L., Dhanaraj A.L., Marian C.O., Ehlenfeldt M.K. & Vinyard B. (2004). Dehardening kinetics, bud development, and dehydrin metabolism in blueberry cultivars during deacclimation at constant, warm temperatures. Journal of the American Society for Horticultural Sciences 129, 667–674.

Berry J. & Björkman O. (1980). Photosynthetic response and adaptation to temperature in higher plants. Annual Review of Plant Physiology 31, 491–543.

Bigg E.K. (1953). The supercooling of water. Proceedings of the Physical Society Section B 66, 688–694.

Bokhorst S.F., Bjerke J.W., Tømmervik H., Callaghan T.V. & Phoenix G.K. (2009). Winter warming events damage sub-Arctic vegetation: consistent evidence from an experimental manipulation and a natural event. Journal of Ecology 97, 1408–1415.

Burke M.J., Gusta L.V., Quamme H.A., Weiser C.J. & Li P.H. (1976). Freezing and injury in plants. Annual Review of Plant Physiology 27, 507–528.

Cannell M.G.R. & Smith R.I. (1983). Thermal time, chill days and prediction of budburst in *Picea sitchensis*. Journal of Applied Ecology 20, 951–963.

Cannell M.G.R. & Willett S.C. (1975). Rates and times at which needles are initiated in buds of different provenances of *Pinus contorta* and *Picea sitchensis* in Scotland. Canadian Journal of Forest Research 5, 367–380.

Chmielewski F., Götz K., Weber K.C. & Moryson S. (2018). Climate change and spring frost damages for sweet cherries in Germany. International Journal of Biometeorology 62, 217–228.

Citadin I., Raseira M.C.B., Herter F.G. & Silva J.B. (2001). Heat requirement for blooming and leafing in peach. HortScience 36, 305–307.

Cleland E.E., Chuine I., Menzel A., Mooney H.A. & Schwartz M.D. (2007). Shifting plant phenology in response to global change. Trends in Ecology & Evolution 22, 357–365.

Cook C. & Jacobs G. (2000). Progression of apple (*Malus × domestica* Borkh.) bud dormancy in two mild winter climates. The Journal of Horticultural Science and Biotechnology 75, 233–236.

Couvillon G.A. & Erez A. (1985). Influence of prolonged exposure to chilling temperatures on bud break and heat requirement for bloom of several fruit species. Journal of the American Society for Horticultural Sciences 110, 47–50.

Cragin J., Serpe M., Keller M. & Shellie K. (2017). Dormancy and cold hardiness transitions in winegrape cultivars Chardonnay and Cabernet Sauvignon. American Journal of Enology and Viticulture 68, 195–202.

Dantec C.F., Vitasse Y., Bonhomme M., Louvet J., Kremer A. & Delzon S. (2014). Chilling and heat requirements for leaf unfolding in European beech and sessile oak populations at the southern limit of their distribution range. International Journal of Biometeorology 58, 1853–1864.

Dry P. & Coombe B. (2004). Grapevine growth stages-The modified EL system. Viticulture 1-Resources.

Fan S., Bielenberg D.G., Zhebentyayeva T.N., Reighard G.L., Okie W.R., Holland D. & Abbott A.G. (2010). Mapping quantitative trait loci associated with chilling requirement, heat requirement and bloom date in peach (*Prunus persica*). New Phytologist 185, 917–930.

Farokhzad A., Nobakht S., Alahveran A., Sarkhosh A. & Mohseniazar M. (2018). Biochemical changes in terminal buds of three different walnut (*Juglans regia* L.) genotypes during dormancy break. Biochemical Systematics and Ecology 76, 52–57.

Farquhar G.D. & Richards R.A. (1984). Isotopic composition of plant carbon correlates with water-use efficiency of wheat genotypes. Functional Plant Biology 11, 539–552.

Ferguson J.C., Tarara J.M., Mills L.J., Grove G.G. & Keller M. (2011). Dynamic thermal time model of cold hardiness for dormant grapevine buds. Annals of Botany 107, 389–396.

Ferguson J.C., Moyer M.M., Mills L.J., Hoogenboom G. & Keller M. (2014). Modeling dormant bud cold hardiness and budbreak in twenty-three Vitis genotypes reveals variation by region of origin. American Journal of Enology and Viticulture 65, 59–71.

Gray L.K. & Hamann A. (2013). Tracking suitable habitat for tree populations under climate change in western North America. Climatic Change 117, 289–303.

Gu L., Hanson P.J., Post W.M., Kaiser D.P., Yang B., Nemani R., …, Meyers T. (2008). The 2007 Eastern US spring freeze: increased cold damage in a warming world? BioScience 58, 253–262.

Horvath D.P., Anderson J.V., Chao W.S. & Foley M.E. (2003). Knowing when to grow: signals regulating bud dormancy. Trends in Plant Science 8, 534–540.

Huang X., Lakso A.N. & Eissenstat D.M. (2005). Interactive effects of soil temperature and moisture on Concord grape root respiration. Journal of Experimental Botany 56, 2651–2660.

Hunter A.F. & Lechowicz M.J. (1992). Predicting the timing of budburst in temperate trees. Journal of Applied Ecology 29, 597–604.

Kalberer S.R., Wisniewski M. & Arora R. (2006). Deacclimation and reacclimation of cold-hardy plants: Current understanding and emerging concepts. Plant Science 171, 3–16.

Kalberer S.R., Leyva-Estrada N., Krebs S.L. & Arora R. (2007). Frost dehardening and rehardening of floral buds of deciduous azaleas are influenced by genotypic biogeography. Environmental and Experimental Botany 59, 264–275.

Kolstad E.W., Breiteig T. & Scaife A.A. (2010). The association between stratospheric weak polar vortex events and cold air outbreaks in the Northern Hemisphere. Quarterly Journal of the Royal Meteorological Society 136, 886–893.

Lang G.A., Early J.D., Martin G.C. & Darnell R.L. (1987). Endo-, para-, and ecodormancy: physiological terminology and classification for dormancy research. HortScience 22, 371–377.

Lloyd J. & Firth D. (1990). Effect of defoliation time on depth of dormancy and bloom time for low-chill peaches. HortScience 25, 1575–1578.

Londo J.P. & Johnson L.M. (2014). Variation in the chilling requirement and budburst rate of wild Vitis species. Environmental and Experimental Botany 106, 138–147.

Londo J.P. & Kovaleski A.P. (2017). Characterization of wild North American grapevine cold hardiness using differential thermal analysis. American Journal of Enology and Viticulture 68, 203–212.

Luedeling E., Girvetz E.H., Semenov M.A. & Brown P.H. (2011). Climate change affects winter chill for temperate fruit and nut trees. PLoS ONE doi.org/10.1371/journal.pone.0020155

Mills L.J., Ferguson J.C. & Keller M. (2006). Cold-hardiness evaluation of grapevine buds and cane tissues. American Journal of Enology and Viticulture 57, 194–200.

Pagter M. & Arora R. (2013). Winter survival and deacclimation of perennials under warming climate: physiological perspectives. Physiologia Plantarum 147, 75–87.

Penfield S. (2008). Temperature perception and signal transduction in plants. New Phytologist 179, 615–628.

Pérez F.J. & Burgos B. (2004). Alterations in the pattern of peroxidase isoenzymes and transient increases in its activity and in H_2_O_2_ levels take place during the dormancy cycle of grapevine buds: the effect of hydrogen cyanamide. Plant Growth Regulation 43, 213–220.

Pérez F.J. & Lira W. (2005). Possible role of catalase in post-dormancy budbreak in grapevines. Journal of Plant Physiology 162, 301–308.

Rigano V.D.M., Vona V., Lobosco O., Carillo P., Lunn J.E., Carfagna S., …, Rigano C. (2006). Temperature dependence of nitrate reductase in the psycrophilic unicellular alga *Koliella antarctica* and the mesophilic alga *Chlorella sorokiniana*. Plant, Cell and Environment 29, 1400–1409.

Rubio S., Dantas D., Bressan-Smith R. & Pérez F.J. (2016). Relationship between endodormancy and cold hardiness in grapevine buds. Journal of Plant Growth Regulation 35, 266–275.

Ruelland E. & Zachowski A. (2010). How plants sense temperature. Environmental and Experimental Botany 69, 225–232.

Salazar-Gutiérrez M.R., Chaves B., Anothai J., Whiting M. & Hoogenboom G. (2014). Variation in cold hardiness of sweet cherry flower buds through different phenological stages. Scientia Horticulturae 172, 161–167.

Salazar-Gutiérrez M.R., Chaves B. & Hoogenboom G. (2016). Freezing tolerance of apple flower buds. Scientia Horticulturae 198, 344–351.

Shaltout A.D. & Unrath C.R. (1983). Rest completion prediction model for ‘Starkrimson Delicious’ apples. Journal of the American Society for Horticultural Sciences 108, 957–961.

Szalay L., Molnár A. & Kovács S. (2017). Frost hardiness of flower buds of three plum (*Prunus domestica* L.) cultivars. Scientia Horticulturae 214, 228–232.

Takeuchi M. & Kasuga J. (2017). Bark cells and xylem cells in Japanese white birch twigs initiate deacclimation at different temperatures. Cryobiology doi.org/10.1016/j.cryobiol.2017.11.007

Taulavuori K.M.J., Taulavuori E.B., Skre O., Nilsen J., Igeland B. & Laine K.M. (2004). Dehardening of mountain birch (*Betula pubescens* ssp. *czerepanovii*) ecotypes at elevated winter temperatures. New Phytologist 162, 427–436.

Truhlar D.G. & Kohen A. (2001). Convex Arrhenius plots and their interpretation. Proceedings of the National Academy of Sciences 98, 848–851.

Vitasse Y., Lenz A. & Körner C. (2014). The interaction between freezing tolerance and phenology in temperate deciduous trees. Frontiers in Plant Science doi: 10.3389/fpls.2014.00541

Walsh J., Wuebbles D., Hayhoe K., Kossin J., Kunken K., Stephens G., …, Somerville R. (2014). Chapter 2: Our Changing Climate. In Climate Change Impacts in the United States: The Third National Climate Assessment (eds J.M. Melillo, T. Richmond & G.W. Yohe), pp. 19–67. U.S. Global Change Research Program doi: 10.7930/J0KW5CXT

Weinbaum S.A., Polito V.S. & Muraoka T.T. (1989). Assessment of rest completion and its relationship to appearance of tetrads in anthers of ‘Nonpareil’ almond. Scientia Horticulturae 38, 69–76.

Wolf T.K. & Cook M.K. (1992). Seasonal deacclimation patterns of three grape cultivars at constant, warm temperature. American Journal of Enology and Viticulture 43, 171–179.

Wolf T.K. & Cook M.K. (1994). Cold hardiness of dormant buds of grape cultivars: comparison of thermal analysis and field survival. HortScience 29, 1453–1455.

Zhang J. & Taylor C. (2011). The dynamic model provides the best description of the chill process on ‘Sirora’ pistachio trees in Australia. HortScience 46, 420–425.

